# Novel Approach for Refractive Index Matching in Calcified Tissues

**DOI:** 10.1101/2024.07.23.604611

**Authors:** Damien Laudier, Macy Mora-Antionette, Andi Garcia-Ortiz, Karl J. Lewis

## Abstract

Tissue-clearing techniques have become indispensable in biomedical research, especially for three-dimensional (3D) imaging. These techniques enable the visualization of complex biological structures by rendering tissues transparent while preserving their structural integrity. Calcified tissues, such as bone and cartilage, pose a unique challenge in imaging studies due to their inherent opacity and rigidity. In this work, we introduce a novel modification to the BABB clearing protocol aimed at enhancing the clearing efficacy specifically for calcified tissues. Through this work, we provide an enhanced tissue-clearing method that addresses the specific challenges associated with calcified tissues. This advancement has the potential to facilitate more precise and comprehensive studies in fields such as developmental biology, orthopedics, and regenerative medicine.

## Introduction

Tissue-clearing techniques have become indispensable in biomedical research, especially for three-dimensional (3D) imaging[1–4]. These techniques enable the visualization of complex biological structures by rendering tissues transparent while preserving their structural integrity. One of the classical methods employed in tissue clearing is the Benzyl Alcohol and Benzyl Benzoate (BABB) technique, which has been widely utilized due to its simplicity and effectiveness[2,5,6]. However, despite its broad applicability, the BABB method faces significant challenges when applied to calcified tissues, notorious for their opacity and density.

Calcified tissues, such as bone and cartilage, pose a unique challenge in imaging studies due to their inherent opacity and rigidity. Traditional clearing techniques often fall short in these tissues, limiting the resolution and depth of imaging[7]. Therefore, optimizing tissue-clearing protocols for calcified tissues is critical for advancing our understanding of bone biology, pathology, and regeneration.

In this work, we introduce a novel modification to the BABB clearing protocol aimed at enhancing the clearing efficacy specifically for calcified tissues. Our modified protocol leverages adjustments in the composition and application of clearing agents to improve the transparency and preservation of calcified tissues. By systematically optimizing each step of the process, we aim to achieve superior clearing while maintaining the structural and molecular integrity of the tissues.

This paper details our modified BABB clearing protocol. We present a comprehensive analysis of its efficacy compared to the traditional BABB method and other existing techniques. Furthermore, we demonstrate the applicability of our protocol in light-sheet microscopy. Our results show significant improvements in the visualization of calcified tissues, opening new avenues for detailed anatomical and functional studies.

Through this work, we provide an enhanced tissue-clearing method that addresses the specific challenges associated with calcified tissues. This advancement has the potential to facilitate more precise and comprehensive studies in fields such as developmental biology, orthopedics, and regenerative medicine.

## Methods

### Reagents

- Z-FIX/Zinc Formalin: NC9937162 (Fisher Scientific)
- EDTA: E5134-1KG (Sigma-Aldrich)
- 30% Hydrogen Peroxide: 470301-282 (VWR)
- Acetonitrile: A998-1 (Fisher Scientific)
- Ethylene Glycol Monoethyl Ether/2-Ethoxyethanol: E180-1 (Fisher Scientific)
- Benzyl Alcohol: 402834-1L (Sigma Aldrich)
- Benzyl Benzoate: B6630 (Sigma-Aldrich)
- PBS with Tween 20 (PBST; Sigma-Aldrich)

### Equipment

- 15 mL conical tubes
- 50 mL conical tubes
- Eppendorf tubes
- Weighing boats
- Electronic scale
- Disposable plastic spatulas
- Hot plate with stirrer
- Shaker with temperature control
- pH meter
- 10 μL micropipette
- 100 μL micropipette
- 1000 μL micropipette
- 5 mL electronic pipette
- 50 mL electronic pipette

#### Decalcified Protocol

1. Perfuse mice with 10% Z-FIX. Dissect bones and place them in Z-FIX overnight at 4°C.
  - Technical note: If perfusion is not possible, fixation time should be increased accordingly.
2. Decalcify bones in 20% EDTA, pH 7-7.5 at 37°C for 5 days. Change solutions every day. Rinse samples well with DI water before the next step.
  - **Technical note**: Other decalcification reagents may be used. If using EDTA, decalcification can take longer depending on the reagent source and sample dimensions. As such, decalcification should be verified using Micro-CT analysis. Zinc-based fixatives may cause salt artifacts in Micro-CT images and need to be thresholded out.
3. Dilute the 30% stock hydrogen peroxide in DI water by 2x. Incubate samples in 15% hydrogen peroxide for 20 minutes or until marrow becomes opaque, then rinse well with DI water.
4. Whole-mount immunofluorescent staining of samples takes place at room temperature with constant shaking. You can use 15 mL conical tubes and Eppendorf tubes for this part. Each step takes 24 hours to complete. Starting at the secondary antibody, cover all your tubes with aluminum foil for the rest of the process to avoid.
  - Block using 10% BSA in PBST overnight. If you know the species of the primary antibodies, feel free to instead use 3% BSA with 10% species serum.
  - Dilute 1:100 primary antibody in blocking buffer.
  - Wash in PBST.
  - Dilute 1:200 secondary antibody in PBST.
  - Wash in PBST.
5. Incubate samples in 2-ethoxyethanol for 4 hours to overnight.
6. Pour off 2-ethoxyethanol and replace with acetonitrile with three changes over 4 hours to overnight.
7. Pour off acetonitrile and replace with benzyl alcohol for 2-4 hours.
8. Pour off benzyl alcohol and replace it with benzyl benzoate. Incubate overnight until samples are fully transparent. Samples may be stored in benzyl benzoate until ready for histology processing or imaging. If imaging, be careful as benzyl benzoate is an organic solvent that can damage many objectives.

#### Undecalcified Protocol

1. 1. Perfuse mice with 10% Z-FIX. Dissect bones and place them in Z-FIX overnight at 4°C.
  - Technical note: If perfusion is not possible, fixation time should be increased accordingly.
2. Post-fixation: rinse samples well with DI water.
3. Incubate samples in 30% hydrogen peroxide for 90 minutes or until marrow becomes opaque, then rinse well with DI water.
4. Whole-mount immunofluorescent staining of samples takes place at room temperature with constant shaking. You can use 15 mL conical tubes and Eppendorf tubes for this part. Each step takes 24 hours to complete. Starting at the secondary antibody, cover all your tubes with aluminum foil for the rest of the process to avoid.
  - Block using 10% BSA in PBST overnight. If you know the species of the primary antibodies, feel free to instead use 3% BSA with 10% species serum.
  - Dilute 1:100 primary antibody in blocking buffer.
  - Wash in PBST.
  - Dilute 1:200 secondary antibody in PBST.
  - Wash in PBST.
5. Incubate samples in 2-ethoxyethanol for 8 hours to overnight.
6. Pour off 2-ethoxyethanol and replace with acetonitrile with three changes over 24 hours.
7. Pour off acetonitrile and replace with benzyl alcohol for 8 hours to overnight.
8. Pour off benzyl alcohol and replace with benzyl benzoate. Incubate until samples are fully transparent (time may vary depending on bone size).
9. Samples may be stored in benzyl benzoate until ready for histology processing.

## Results

Our approach provided successful clearing of bone tissue. To assess the ability to perform bulk immunohistochemistry and light sheet imaging, we processed our samples through a protocol for labeling nerves in bone tissue. We labeled tyrosine hydroxylase (TH) and vesicular acetylcholine transferase (VAChT) which represent adrenergic and cholinergic nerves, respectively. Compared to the current state-of-the-art protocol for clearing mineralized tissues, PEGASOS, our bone was clearer (Fig 1 A-B). The availability of binding epitopes for antibodies was confirmed via chromogenic stain of plastic-embedded sections of cleared bone tissue (Fig 1 C-F).

**Figure 1:**
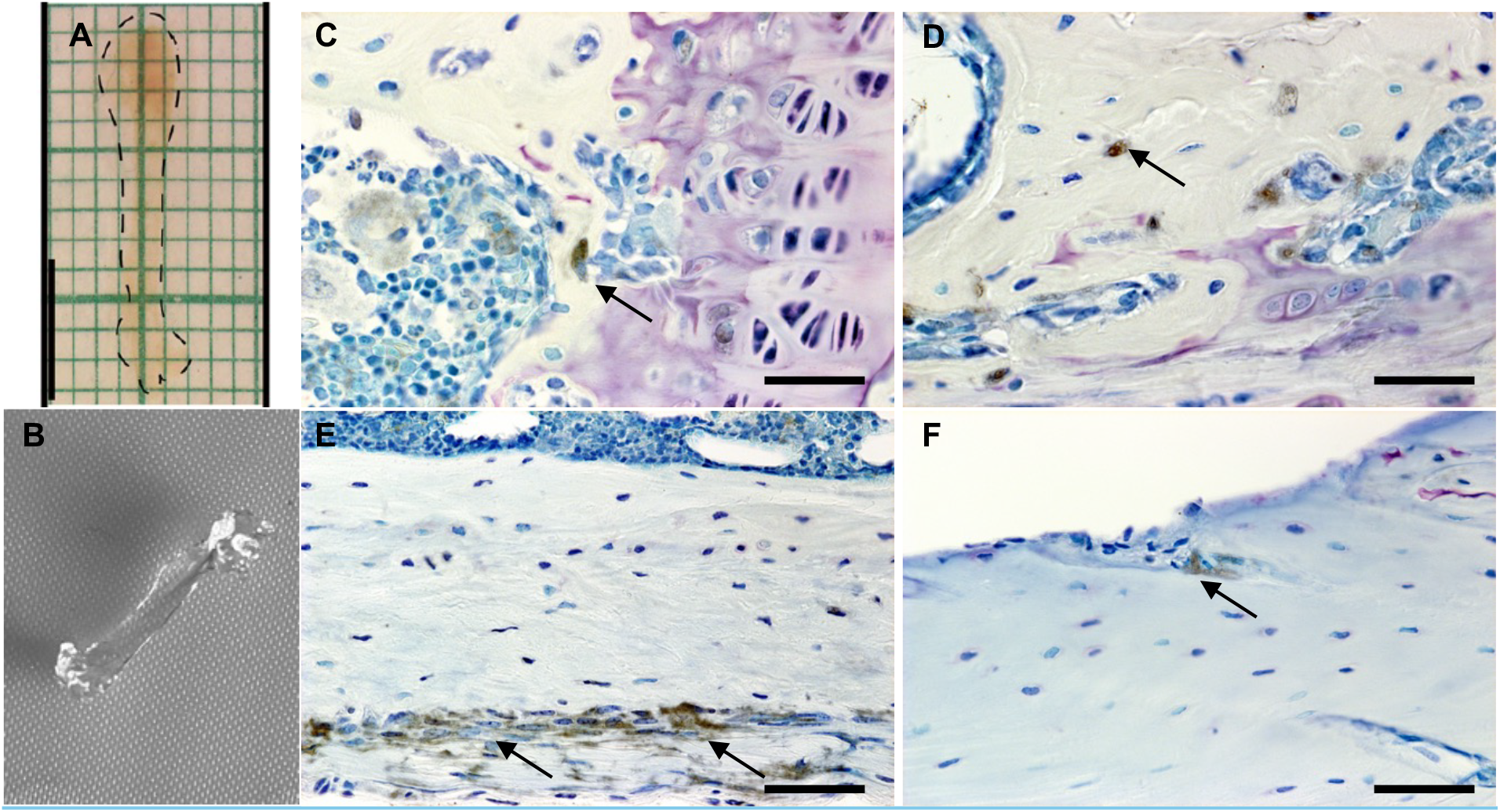
A) PEGASOS cleared bone and B) bone cleared by our modified approach. Both samples are adult mouse femurs. Note that our new approach removes more of the marrow space pigment, offering the potential for higher quality resolution of fluorescent markers via 3D imaging of intact samples. C-F, Plastic embedded sections of clear bone. Chromogenic stain was used to label VAchT for cholinergic fibers with a toluidine blue counterstain (scale bar 100μm). Cholinergic fibers are shown to permeate trabecular (panel A) and cortical bone (panels B-D), as indicated by the black arrows. These images confirm that our clearing method does not damage binding epitopes and the presence of cholinergic fibers in bone matrix near embedded osteocytes. (PEGASOS image sourced from D. Jing et al, Cell Research 2018)

Bulk immunohistochemistry for TH (red) and VAChT (green) was successful. In the example images here, we show a mouse third metatarsal (MT3) that has been whole-mount labeled and imaged using light sheet microscopy (Fig 2). Note that nerves of both types can be seen to track together (indicated by a yellow color) and individually, and that the patterns of their anatomy differ depending on location along the length of the bone. Moreover, extensively labeled features can be seen in the marrow space. These results exhibit the utility of our approach and the opportunity for analyses of biological/anatomical features in intact bone tissues.

**Figure 2:**
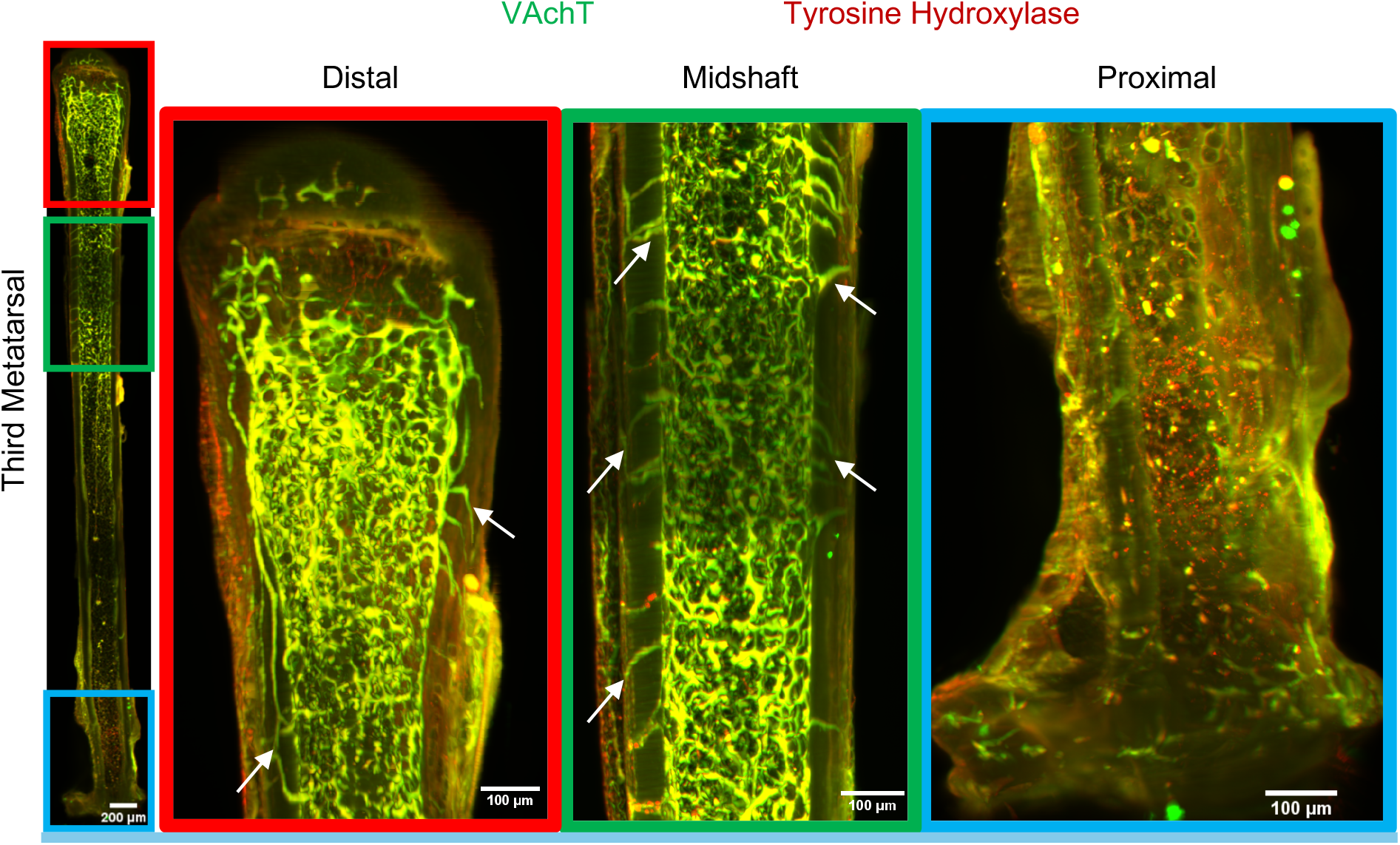
3D projection light sheet microscopy images of a cleared bones bulk immunohistochemistry-stained mouse third metatarsal (MT3) with VAchT (Green) and tyrosine hydroxylase (Red) representing cholinergic and adrenergic nerve fibers, respectively. At the distal and proximal ends of the bone clear distinct nerve labeling can be seen. At the midshaft, transcortical innervation can be observed, with apparent extensive branching throughout the distal half of the bone. White arrows indicate transcortical cholinergic fibers. These preliminary results validate our approach and showcase the ability to map nerves in 3-dimensions.

## Discussion

Tissue-clearing is an indispensable tool for three-dimensional imaging. Through adjustments in the composition and application of clearing agents, our novel method of tissue-clearing addresses the issues faced by traditional techniques, providing enhanced transparency and structural integrity for calcified tissues. In turn, we can achieve higher image resolution and imaging depth for our samples.

Our cleared tissue samples can be employed in a plethora of imaging modalities. In this paper, we highlighted the enhanced efficacy of our cleared samples during confocal and light-sheet microscopy. The results confirmed our method’s superiority to traditional BABB clearing in calcified tissue visualization and structural integrity.

Through this advancement, researchers can now facilitate more precise and comprehensive studies in fields, such as developmental biology, orthopedics, and regenerative medicine. For bone in particular, our tissue-clearing efforts have allowed us to interrogate questions surrounding nerve innervations within long bones. Our cleared samples are also not limited to confocal and light-sheet microscopy, other imaging modalities such as two-photon and three-photon microscopy can also be implemented. The potential for advanced tissue-clearing is immense and has created a solution to historical barriers, due to the opacity and density of calcified tissues, and will be central to novel discoveries moving forward.

## Notes

### Competing Interest Statement

The authors have declared no competing interest.

### Summary of Updates

In error, we neglected to include a fourth author. That author has been added here.

